# In garden dormouse cerebral cortex, specific transcriptional programs exist for all major phases of hibernation

**DOI:** 10.64898/2026.03.10.710569

**Authors:** Adrian Jakubowski-Addabbo, Maarten R. Hamberg, Fia Fia Lie, James Gray, Roelof A. Hut, Maurits Roorda, Victor Guryev, Rob H. Henning

**Affiliations:** Department of Clinical Pharmacy and Pharmacology, University Medical Center Groningen, University of Groningen, the Netherlands; Chronobiology Unit, Groningen Institute for Evolutionary Life Sciences, University of Groningen, Groningen, 9747 AG the Netherlands; European Research Institute for the Biology of Ageing, University Medical Center Groningen, University of Groningen, Groningen, The Netherlands; Department of Pharmacology, Medical Faculty, Universitas Tarumanagara, Jakarta, Indonesia

**Keywords:** hibernation, cerebral cortex, RNA-seq, torpor–arousal cycle, metabolic adaptation, neuroprotection

## Abstract

Hibernators cycle between torpor, a state of profound metabolic and thermoregulatory suppression, and brief arousals during which metabolic rate and body temperature rapidly return to euthermic levels. These repeated physiological pressures require robust mechanisms to preserve brain integrity. Because the cerebral cortex is not thought to control hibernation directly yet must remain viable throughout torpor and recover rapidly during arousal, it provides a useful model for studying neural adaptation to hibernation. We therefore performed RNA sequencing of cerebral cortex from garden dormice (*Eliomys quercinus*) sampled during summer euthermia (SE), early torpor (TE), late torpor (TL), early arousal (AE), and late arousal (AL).

Differential expression analysis revealed strongly stage-specific transcriptional remodeling across the hibernation cycle. Entry into torpor (SE–TE) and the transition from early to late arousal (AE–AL) showed minimal change, with 16 and 2 differentially expressed genes (DEGs), respectively. In contrast, extensive regulation was observed during torpor progression (TE–TL; 576 DEGs) and especially during the transition from late torpor to early arousal (TL–AE; 697 DEGs). Intermediate numbers of DEGs were detected in AL–TE (260) and AL–SE (50). Principal component and enrichment analyses indicated that the dominant axes of variation were associated with RNA processing and proteostatic control, metabolic and redox-related adaptation, and changes in intracellular trafficking and protein handling. In addition, comparison of adjacent contrasts revealed a marked opposite-direction transcriptional reversal between TE–TL and TL–AE, consistent with coordinated reactivation of torpor-associated programs during arousal.

Together, these findings support a model in which cortex adaptation to hibernation involves transcriptional reprogramming consistent with metabolic suppression during torpor progression, especially in pathways related to carbohydrate and central carbon metabolism, redox homeostasis, and cellular signalling, followed by rapid reversal of these programs during early arousal.

## Introduction

Hibernators have physiological adaptations enabling them to survive extreme environmental conditions. Most important among these is the ability to enter torpor, a state characterized by a sharp reduction in metabolic activity, allowing animals to conserve energy and endure periods of resource scarcity(Geiser, 2004). During deep torpor, metabolic rate can decline to as low as 2–5% of basal level, resulting in body temperature (Tb) aligning closely with ambient temperatures (Ta). For example, the seasonal hibernator garden dormouse (*Eliomys quercinus*), has torpor bouts lasting up to 14 days with body temperature decreasing to ∼2 °C above Ta(Giroud et al., 2020; Pajunen, 1983). Its torpor bouts are alternated with short interbout arousals (iBAs), typically lasting about half a day, during which normal metabolic rate and euthermic temperature are rapidly restored(Ruf et al., 2021). While iBAs are present in all hibernators, their function is still elusive, although they are believed to allow for physiological repair and recovery from torpor-induced damage(Arendt et al., 2003; de Wit et al., 2023; Talaei et al., 2012).

At the molecular level, torpor–arousal cycles imply reversible control of energy use and cellular maintenance, as programs suppressed during torpor must be reactivated quickly at arousal, while damage-limiting and repair pathways may be selectively engaged at specific transition points. This makes the timing of transcriptional and post-transcriptional regulation a central question when interpreting brain adaptations across torpor and iBA.

A key constraint in the brain is that deep torpor combines (i) low temperature, which slows enzymatic reaction rates, (ii) reduced ATP turnover and cerebral energy demand(Henry et al., 2007), and (iii) a need to preserve neuronal membranes, synapses, and mitochondria for rapid functional recovery. This creates a logic of *gated maintenance*: during torpor progression, energy-expensive programs (biosynthesis, signaling, immune activation) should be suppressed, while protective programs (redox control, mitochondrial quality control, damage containment) are maintained. During arousal onset, these programs must flip quickly to support restoration of ionic homeostasis, synaptic function, and network activity, consistent with the intense NREM rebound observed early in arousal.

With the advent of more affordable sequencing techniques, RNA-Seq has been increasingly used to investigate physiological adaptations in several hibernating species. While most transcriptomic studies primarily focused on peripheral tissues(Coussement et al., 2023; Hampton et al., 2013; Jansen et al., 2019), understanding brain-specific adaptations is essential, as the brain is likely to orchestrate essential elements of hibernation(Hrvatin et al., 2020; Takahashi et al., 2020) and is particularly vulnerable to even short episodes of hypoxia or nutrient deprivation. Importantly, the brain is among the earliest organs to rewarm, rapidly regaining function during iBA, primarily driven by the activation of perivascular brown fat tissue. Investigating the brain thus provides valuable insights into the mechanisms that preserve neural integrity and function under conditions of metabolic suppression and subsequent rapid rewarming(Carey et al., 2003; Geiser, 2004). Brain RNA-Seq studies conducted used whole brain(Lei et al., 2014; Nespolo et al., 2018) or focused on the hypothalamus, either alone(Haugg et al., 2024) or in relation to other brain areas(Fu et al., 2020; Schwartz et al., 2013). This pattern is supported by hypothalamus transcriptome datasets across hibernation stages, which repeatedly implicate circadian/clock regulation, stress-response and metabolic remodeling as major axes of variation(Morrison et al., 2025; Schwartz et al., 2013). Hypothalamus is a region integrating thermoregulation, energy balance, and circadian and seasonal rhythmicity adjustments, and remains active during torpor and arousal(Sonntag & Arendt, 2019), and as such widely regarded as an important brain region orchestrating hibernation. Between these studies, common molecular pathways that have been implicated as being regulated between torpor and arousal in hypothalamus include extracellular matrix organization, antioxidant defence, circadian rhythm, mRNA surveillance, DNA damage response and stress response. In addition, initiation of fasting-induced torpor in mice hinges on specific hypothalamic regions(Hrvatin et al., 2020; Takahashi et al., 2020). In addition to changes in gene expression, the brain of hibernators undergoes extensive morphological and molecular remodeling during torpor-arousal cycles. Importantly, the torpid brain endures a significant loss in synapse complexity and density(Popov & Bocharova, 1992) and a substantial tau hyperphosphorylation(Arendt et al., 2003), all of which are reversed during iBA.

Despite this progress, two knowledge gaps remain. First, it is still unclear which brain transcriptional programs are conserved across hibernators (and which are species- or tissue-specific), because studies differ in species, brain region, and torpor staging. Second, while the hypothalamus is central for torpor control, less is known about how cortical tissue itself implements protection and recovery across torpor progression and rewarming, i.e., adaptation within neuronal tissue rather than orchestration of torpor onset.

To understand the orchestration of brain adaptations to hibernation, the cortex is an interesting region for transcriptomic analysis. Comparative cortical transcriptomics in other hibernators also supports substantial seasonal remodeling of neuronal tissue, suggesting cortex is not just a passive target but engages in active adaptation programs(Faherty et al., 2016). First, it likely represents adaptation of neuronal tissue to hibernation rather than displaying changes in gene expression related to both controlling and adapting to torpor-arousal cycles such as in hypothalamus. Further, in addition to synaptic remodeling and hyperphosphorylation of tau, the torpid cerebral cortex is subject to drastic, yet reversible, changes in activity. While the cortex is virtually isoelectric during torpor, some hibernator species subsequently display intense NREM sleep during arousals, characterized by low frequency spectral power at the arousal onset(Daan et al., 1991; Trachsel et al., 1991). Torpor is furthermore associated with neuroinflammatory modulation in several brain regions, including hippocampus(Cogut et al., 2018). To chart cortical adaptation across the torpor–arousal cycle, we performed RNA-seq on garden dormouse cerebral cortex during summer euthermia (SE) and four hibernation states spanning early vs late torpor (TE, TL) and early vs late arousal (AE, AL). We first ask when the cortex undergoes major transcriptional remodeling (entry into torpor vs progression within torpor vs arousal onset), and whether regulation is reversible across the TL-AE transition. We then complement gene-level differential expression with pathway-level enrichment to identify coordinated functional programs.

## Methods

### Animals and Tissue Sampling

Experimental protocols adhered to the guidelines of the Animal Welfare Body at the University of Groningen and were authorized by the Central Committee on Animal Experiments of the Netherlands, license AVD1050020198666. Animal care, monitoring, and hibernation induction followed the methods outlined by de Wit et al. (de Wit et al., 2023). Twenty garden dormice (10 males and 10 females) were transferred from outside group housing under natural conditions to laboratory housing, where they were individually kept in plexiglass cages equipped with wooden nest boxes lined with coconut shreds(Hut et al., 2002). Nest box temperature (Tn) was continuously monitored with a HOBO Onset pendant (MX2201) to register torpor bouts and iBA. Animals had unrestricted access to water, sunflower seeds, and Altromin 7024 chow, though hibernating individuals rarely consumed these. Hibernation was induced by placing animals in constant darkness followed by a gradual lowering of ambient temperature (Ta) from 20°C to 5 °C, which was maintained for at least 4 weeks. Details regarding sex, body weight, age, and the duration of torpor and arousal cycles are presented in Table S1.

### Tissue Collection

Animals were placed in a darkened chamber filled with 5% isoflurane in air, transferred to an operating room where rectal temperature was immediately recorded, with subsequent maintenance of anesthesia with isoflurane in air. An abdominal midline incision was made and blood was collected via cardiac puncture, followed by a whole-body perfusion using 150 mL of 0.9% saline solution. Torpid animals were perfused with ice-cold saline and placed on ice, while euthermic animals were perfused at room temperature. Subsequently, brain tissues were harvested, snap-frozen in liquid nitrogen, and stored at -80°C. The study included animals in the following physiological states: summer euthermic (SE, collected in July, n=3), early torpor (TE, approximately 24 hours into torpor, n=5), late torpor (TL, more than 8 days in torpor, n=5), early arousal (AE, 1 hour after spontaneous arousal onset, n=4), and late arousal (AL, 8 hours into spontaneous arousal, n=4).

### Tissue Extraction and Sequencing

RNA was isolated from the brain cortex (NucleoSpin RNA isolation kit, Macherey-Nagel). Libraries were prepared using the NEBNext Ultra II Directional RNA Library Prep Kit, which included ribosomal RNA depletion, directional cDNA synthesis, adapter and Unique Molecular Identifier (UMI) ligation, and amplification. All samples underwent Illumina-based RNA sequencing at a depth of 25x. To reduce amplification duplicates, each read was assigned a unique molecular identifier. Sequencing was performed with 150 bp paired-end reads on an Illumina NextSeq platform, targeting 50 million reads per sample using a stranded sequencing library. Samples yielding fewer than 50 million reads were resequenced to meet the target read count. Library preparation and sequencing were conducted by GenomeScan (Leiden, the Netherlands). Raw forward and reverse reads were delivered in compressed fastq files and stored on the in-house computer cluster of the European Research Institute for the Biology of Ageing (ERIBA, Groningen, the Netherlands).

### RNA-seq read mapping

Raw sequencing data were processed to remove low-quality bases (Phred score ≤ 15), adapter contamination, and reads shorter than 15 bp. Sequencer artifacts, including high-confidence poly-G tails, were removed using fastp v0.20.1 (parameters: -l 50 --threads 20 -i -I -o -O)(Chen et al., 2018). Filtered reads were then aligned to a custom-made garden dormouse reference genome. For the initial mapping, a STAR v2.7.0.10b index was generated using --runThreadN 32 --runMode genomeGenerate with default parameters(Dobin et al., 2013). A second mapping pass was performed using a splice junction-enriched STAR index (--runMode genomeGenerate --runThreadN 32 --sjdbFileChrStartEnd --limitSjdbInsertNsj 1500000), which improved splice-site detection(Veeneman et al., 2016). Following alignment, reads were filtered based on unique molecular identifiers (UMIs); only uniquely mapped reads per genomic position were retained, and duplicate reads were removed on a first-come, first-served basis.

### Transcript annotation

Using the RNA-based genome annotation for the garden dormouse, transcript sequences were extracted and compared against Mus musculus reference datasets using BLASTN and BLASTX to assign putative orthologs and standardized gene annotations(McGinnis & Madden, 2004).

### Gene quantification and differential expression analysis

Gene expression was quantified using featureCounts version 2.12.2 from the Rsubread package with the following parameters: annot.ext = genome.annotation, isGTFAnnotationFile = TRUE, strandSpecific = 1, primaryOnly = TRUE, and requireBothEndsMapped = TRUE(Liao et al., 2019). The resulting count matrix was imported into edgeR(Robinson et al., 2010), where gene expression was modeled using a negative binomial distribution. RNA-seq count data were normalized using the trimmed mean of M-values (TMM) method implemented in calcNormFactors(). A design matrix without an intercept was constructed to model group-specific differences, and dispersion estimates were calculated before fitting a generalized linear model with a quasi-likelihood framework. Genes were considered significantly differentially expressed at FDR < 0.01. Specific pairwise comparisons between experimental groups were defined using contrast matrices, and differential expression was assessed for each contrast using quasi-likelihood F-tests. Gene-level results were subsequently merged with the annotation dataset to assign standardized gene names and additional annotation details.

### Visualization

Data visualization was performed using the R packages ggsci, ggplot2, pheatmap, cowplot, enrichplot, scales, VennDiagram, ggrepel, and gridExtra. Log-transformed counts per million (logCPM) of DEGs were extracted for visualization.

### Protein Coding DEGs Analysis

DEGs (FDR < 0.01) were extracted for each contrast, unique gene identifiers, and species-specific IDs (e.g., “ENSSTOG” for ground squirrel) separated from others. Gene biotypes were retrieved via biomaRt by querying the appropriate Ensembl datasets (“itridecemlineatus_gene_ensembl” for squirrel IDs and “mmusculus_gene_ensembl” for mouse IDs). Missing annotations were assigned “NA.” DEGs were then classified as “Protein coding,” “Non protein coding,” or “Not Annotated,” and per-contrast counts and percentages of each category were calculated.

### Principal Component and Functional Enrichment Analyses

To examine the major axes of sample-to-sample variation within the hibernation cycle, we performed principal component analysis (PCA) on normalized gene expression values across hibernation samples. SE samples were excluded because they represent a seasonal reference state outside the torpor–arousal cycle, whereas the aim of this analysis was to resolve transcriptional structure among hibernation states only. To interpret the biological processes underlying the principal components, we then examined the gene loadings for PC1 and PC2 and selected the top 10% of genes with the largest absolute loading values for each component. These gene sets were subjected to functional enrichment analysis to identify overrepresented GO terms and KEGG pathways using mouse-based annotation resources.

### Opposite-Direction Differential Expression Analysis

To investigate whether transcriptional changes occurring during one transition were reversed during the next, differential expression results from adjacent physiological-state comparisons were systematically compared (SE–TE vs TE–TL, TE–TL vs TL–AE, TL–AE vs AE–AL, AE–AL vs AL–TE, and AL–TE vs AL–SE). Genes significant in one contrast were tested for enrichment among genes significantly regulated in the opposite direction in the following contrast. Analyses were performed for total opposite-direction overlap as well as for specific down-to-up and up-to-down transitions. Statistical significance was assessed relative to the shared background of genes represented across all contrasts, and multiple-testing correction was applied.

## Results

Bulk RNA-Seq was performed on cerebral cortex of garden dormouse during summer euthermia (SE), early and late torpor (TE, TL) and early and late arousal (AE, AL). Summer euthermia samples were obtained outside of the hibernating season (July), while early and late torpor samples were obtained during winter after 24 hours and 8 days of torpor respectively, and early and late arousal samples were obtained after 1 hour and 8 hours of arousal (Fig. 1A). Body temperatures at termination confirmed animals were in the correct phases (Table S1).

**Figure 1.**
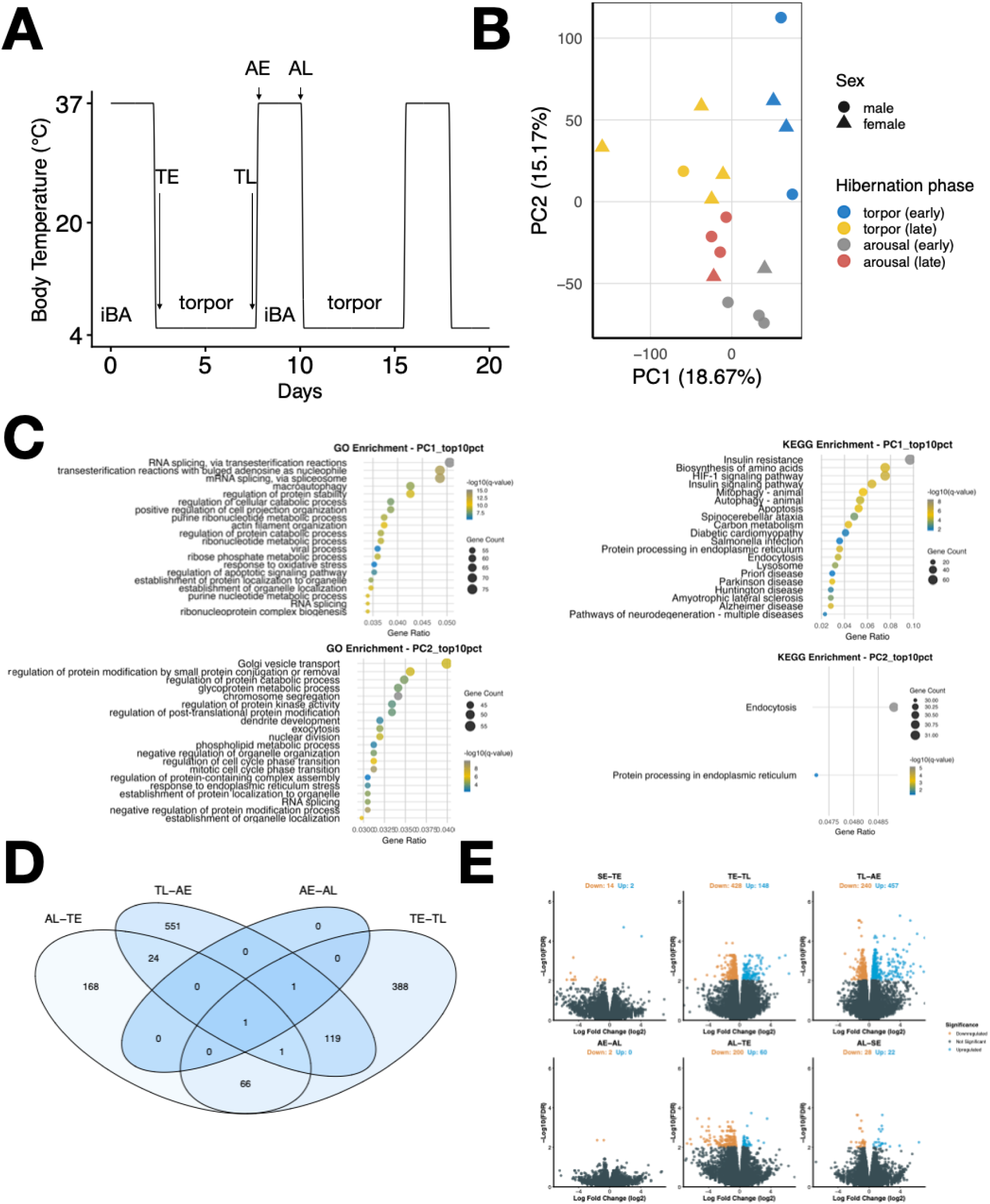
Distinct transcriptional programs characterize major hibernation stages in the garden dormouse cerebral cortex. **(A)** Diagram of timing of sampling during Torpor-Arousal Cycles. In addition to summer euthermic (SE) animals sampled in July (not shown), the diagram depicts the sampling points across distinct hibernation stages comprising early torpor (TE, approximately 24 h into torpor, n=5), late torpor (TL, more than 8 days in torpor, n=5), early arousal (AE, 1 h after spontaneous arousal onset, n=4), and late arousal (AL, 8 hours into spontaneous arousal onset, n=4) **(B)** Sample loading onto principal components 1 and 2. **(C)** GO & KEGG pathway analysis of the top 10% genes driving PC1 & PC2. **(D)** Venn diagram showing overlap and contrast-specific DEGs (FDR < 0.01) across pairwise comparisons between hibernation states (e.g., TE vs TL, TL vs AE, AE vs AL, AL vs TE). **(E)** Volcano plots of DEGs in six contrasts between hibernation stages.

We present the results in three steps. First, we assess global structure across hibernation states using PCA. Second, we quantify differential expression across six biologically motivated contrasts spanning seasonal and within-cycle transitions. Third, we test for reversal of regulation across adjacent transitions (especially TE→TL vs TL→AE).

To obtain an overview of global transcriptional relationships across hibernation stages, we performed PCA on normalized gene-expression values excluding SE samples (Fig. 1B). PC1 separated TE and TL samples, indicating progressive transcriptional remodeling within torpor, whereas PC2 separated torpor from arousal samples, reflecting the major shift associated with rewarming. AE and AL overlapped substantially along PC2, consistent with the minimal differential expression observed between these stages. Together, PC1 and PC2 explained 18.7% and 15.2% of the variance, respectively.

To explore the biological processes contributing to these major axes of variation, we performed GO and KEGG enrichment analysis on the top 10% of genes with the highest absolute loadings on PC1 and PC2 (Fig. 1C). Genes contributing to PC1 were strongly enriched for RNA-processing pathways, particularly RNA splicing, mRNA splicing via the spliceosome, and ribonucleoprotein complex biogenesis. Additional GO terms pointed to macroautophagy, regulation of protein stability and catabolism, purine and ribose phosphate metabolism, actin filament organization, and response to oxidative stress. KEGG analysis supported this pattern by highlighting carbon metabolism, biosynthesis of amino acids, HIF-1 signaling, insulin signaling and insulin resistance, mitophagy, autophagy, apoptosis, endocytosis, lysosome, and protein processing in the endoplasmic reticulum. Together, these results suggest that PC1 captures a coordinated axis of RNA processing, proteostatic control, metabolic reprogramming, and cellular stress adaptation across torpor progression.

Genes contributing to PC2 were enriched for processes related to intracellular trafficking and protein regulation. GO terms included Golgi vesicle transport, exocytosis, glycoprotein and phospholipid metabolic processes, regulation of protein kinase activity, regulation of post-translational protein modification, regulation of protein catabolic processes, protein-containing complex assembly, response to endoplasmic reticulum stress, and establishment of protein localization to organelles. KEGG analysis identified endocytosis and protein processing in the endoplasmic reticulum as the main enriched pathways. These findings indicate that PC2 reflects shifts in vesicle trafficking, membrane-associated processes, and protein handling that distinguish torpor from arousal states.

To investigate the dynamics of gene expression during specific hibernation transitions, we next examined DEG across six distinct hibernation phase contrasts (FDR < 0.01, Fig. 1D & 1E). Two contrasts capture transitions between summer-active and hibernating animals, namely SE–TE and AL–SE. In addition, four contrasts describe transcriptional changes within the hibernation cycle itself: TE–TL, TL–AE, AE–AL, and AL–TE. Across these six contrasts, differential expression ranged from 2 to 697 genes per transition, and in total 1,601 DEG calls were detected when summing across contrasts (Table 1).

**Table 1.** Features of DEGs in 6 defined hibernation phase transitions (‘contrasts’).

|  |  |  |  |  |  |  |
| --- | --- | --- | --- | --- | --- | --- |
| Contrast | SE-TE | TE-TL | TL-AE | AE-AL | AL-TE | AL-SE |
| **Total DEGs** | 16 | 576 | 697 | 2 | 260 | 50 |
| **Up** | 2 | 148 | 457 | 0 | 60 | 22 |
| **Down** | 14 | 428 | 240 | 2 | 200 | 28 |
| **Unique DEGs** | 3 | 384 | 546 | 0 | 159 | 33 |
| **Annotated** | 8<br>(50.0 %) | 484<br>(84.0 %) | 529<br>(75.9 %) | 2<br>(100 %) | 210<br>(80.8 %) | 43<br>(86.0 %) |
| **Not Annotated** | 8<br>(50.0 %) | 92<br>(16.0 %) | 168<br>(24.1 %) | 0<br>(0%) | 50<br>(19.2%) | 7<br>(14.0%) |
| **Protein coding** | 8<br>(50.0 %) | 484<br>(84.0 %) | 529<br>(75.9 %) | 2<br>(100 %) | 210<br>(80.8 %) | 43<br>(86.0 %) |
| **Non protein coding** | 1<br>(6.3 %) | 0<br>(0 %) | 0<br>(0 %) | 0<br>(0 %) | 0<br>(0 %) | 0<br>(0 %) |

Entry into torpor (SE–TE) and interbout arousal (AE–AL) were characterized by minimal transcriptional adjustments, with only 16 and 2 DEGs detected, respectively. In contrast, extensive transcriptional remodeling was observed during late torpor and the transition into arousal. Specifically, the transition from early to late torpor (TE–TL) involved 576 DEGs, the majority of which were downregulated (428 genes), while the transition from late torpor to early arousal (TL–AE) exhibited the highest DEG count overall, with 697 DEGs, predominantly upregulated (457 genes). Additional, albeit more moderate, transcriptional regulation was observed during the transition from late arousal to torpor (AL–TE; 260 DEGs) and from late arousal to summer activity (AL–SE; 50 DEGs) (Table 1).

Sequential analysis of the contrasts revealed that DEG profiles were largely phase-specific. A substantial proportion of DEGs were uniquely differentially expressed in a single contrast, including 3 unique DEGs in SE–TE, 384 in TE–TL, 546 in TL–AE, 159 in AL–TE, and 33 in AL–SE (Table 1). This high degree of contrast-specific regulation underscores the dynamic and tightly regulated nature of transcriptional programs across the hibernation cycle.

For each contrast, DEGs were further classified based on annotation status. Genes assigned a curated gene symbol through 1:1 orthology with the house mouse (*Mus musculus*) were considered annotated, while genes lacking such orthology were classified as unannotated. Across most contrasts, the majority of DEGs were annotated (and correspondingly classified as protein-coding in this table), ranging from 75.9% in TL–AE to 86.0% in AL–SE (Table 1). Notably, several of the larger contrasts (TE–TL, TL–AE, AL–TE, and AL–SE) contained ∼14–24% unannotated genes, whereas AE–AL showed none (Table 1). These unannotated loci may represent dormouse-specific or rapidly evolving genes relevant to cortical hibernation adaptations.

Overall, these results indicate that differential expression during hibernation is dominated by annotated genes (classified as protein-coding in this table), while a smaller fraction of unannotated transcripts may contribute to species-specific adaptations. Importantly, the most pronounced and directional transcriptional reprogramming occurs during late torpor and early arousal, highlighting these phases as critical regulatory checkpoints within the hibernation cycle. Together, differential expression analysis reveals both strongly state-specific and partially overlapping transcriptomic responses across hibernation phases in the garden dormouse cerebral cortex.

We next tested whether genes differentially expressed in one transition were enriched among genes regulated in the opposite direction in the subsequent transition (Table 2). A strong enrichment was observed for the transition from torpor progression to arousal onset: among the 576 DE genes in the TE–TL contrast, 122 showed opposite-direction regulation in the subsequent TL–AE contrast. This overlap was far greater than expected by chance and was dominated by genes shifting from downregulation during torpor progression to upregulation upon arousal. By contrast, the subsequent TL–AE to AE–AL transition showed only a weak opposite-direction signal, and the remaining adjacent comparisons showed little or no enrichment. Together, these findings indicate that large-scale transcriptional reversal is concentrated at the transition from late torpor to early arousal.

**Table 2.** Comparisons of the oppositely regulated genes against subsequent contrasts and against contrasts comparing non-hibernating and torpid animals.

| Comparisons | **SE-TE vs**<br>**TE-TL** | **TE-TL vs**<br>**TL-AE** | **TL-AE vs**<br>**AE-AL** | **AE-AL vs**<br>**TE-TL** | **SE-TE vs**<br>**AL-SE** |
| --- | --- | --- | --- | --- | --- |
| **Number of**<br>**DEGs** | 16 vs 576 | 576 vs 697 | 697 vs 2 | 2 vs 576 | 16 vs 50 |
| **Number of**<br>**shared DEGs** | 2 | 122 | 2 | 2 | 1 |
| **% of**<br>**oppositely**<br>**regulated**<br>**DEGs** | 50 | 100 | 100 | 0 | 100 |

**Figure 2.**
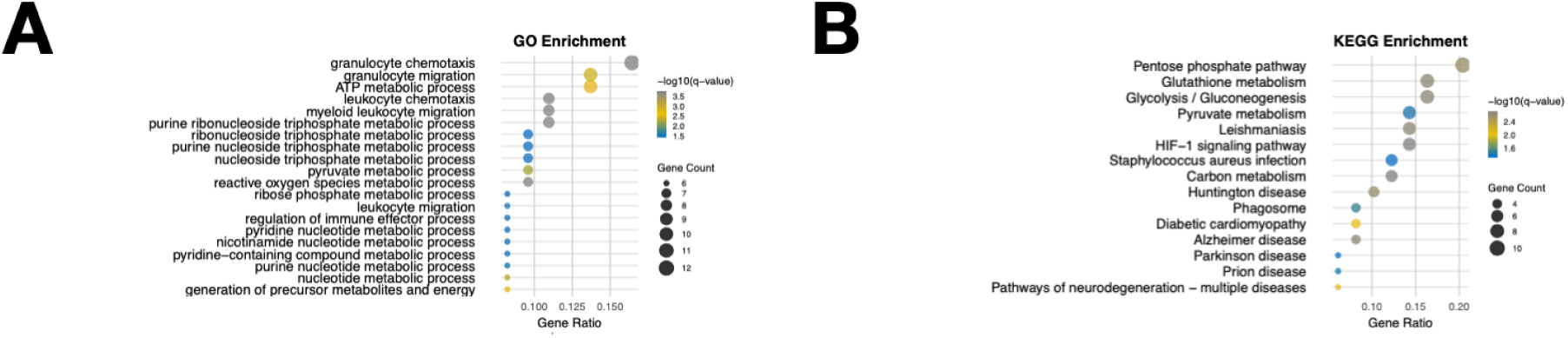
GO & KEGG enrichment of DEGs in hibernation. Depicted are pathways that are oppositely regulated DEGs between the contrasts TE-TL and TL-AE contrasts at FDR < 0.01.

To characterize the biological functions represented in this reversal, we performed pathway enrichment analysis on the oppositely regulated gene set from the TE–TL and TL–AE comparison. KEGG analysis identified a small set of significantly enriched pathways, including pyruvate metabolism as well as disease-labeled pathways such as Parkinson’s disease and diabetic cardiomyopathy, which in this context likely reflect mitochondrial and metabolic stress-response circuitry rather than pathology per se. These results support the interpretation that the torpor-to-arousal transition involves a coordinated reversal of metabolic and cellular maintenance programs.

Using all 26,235 genes shared between contrasts as background, we tested whether DE in one phase were enriched for opposite-direction differential expression in the subsequent phase (FDR < 0.01; genes classified as Up if logFC > 0 and Down otherwise). A strong enrichment was observed for the transition from TL to AE where among the 576 DE genes in the TE–TL contrast, 122 genes showed opposite-direction regulation in the subsequent TL–AE contrast. All 122 of these genes were contained within the TE–TL DEG set, corresponding to a strong enrichment relative to expectation under independence (observed 122 vs expected 2.68; one-sided Fisher’s exact test p = 6.12 × 10⁻²⁰⁹, BH-adjusted q = 3.06 × 10⁻²⁰⁸).

When the two reversal directions tested separately, this enrichment was driven mainly by genes that were downregulated in TE-TL and upregulated in TL-AE (observed 109 vs expected 7.46; p = 3.43 × 10⁻⁹⁶, q = 1.72 × 10⁻⁹⁵). A smaller but still significant group shows the opposite pattern shifting from upregulation in TE-TL to downregulation in TL-AE (13 vs expected 1.35; p = 1.16 × 10⁻⁹, q = 5.81 × 10⁻⁹).

For the subsequent transition from TL–AE to AE–AL, enrichment for opposite-direction regulation was weaker but remained significant (2 observed vs 0.053 expected; p = 7.05 × 10⁻⁴, q = 8.81 × 10⁻⁴), driven exclusively by Up-to-Down regulation. Other adjacent phase comparisons showed minimal or no enrichment for opposite-direction differential expression, indicating that large-scale transcriptional reversals are primarily concentrated at the transition from TL to AE.

## Discussion

Our results show that transcriptional regulation in the garden dormouse cerebral cortex is strongly stage-dependent across the hibernation cycle. Rather than being distributed evenly across all sampled transitions, the most substantial transcriptomic remodeling occurred during torpor progression and, even more prominently, during the transition from late torpor to early arousal. This pattern was evident both in the principal component structure of the dataset and in differential expression analyses, which together identified late torpor and early arousal as the major points of cortical reorganization. In contrast, entry into torpor and the transition from early to late arousal were accompanied by very few differentially expressed genes, indicating that the cortex does not undergo equally large transcriptional shifts at all phases of the torpor–arousal cycle.

The low number of DEGs in SE–TE and AE–AL is biologically informative. One interpretation is that the sampled early torpor time point captures a state in which cortical transcriptional remodeling is still limited, or alternatively that the major adjustments required for torpor entry occur before or outside the sampled window. Similarly, the very small number of DEGs between early and late arousal suggests that a large proportion of transcriptional reactivation may occur very rapidly after arousal onset, with the cortex already approaching a stabilized state by the later arousal time point. In contrast, the high DEG counts in TE–TL and TL–AE indicate that late torpor and arousal onset are the phases during which cortical gene regulation is most dynamic. Together, these findings support a model in which the cortex enters a deeply regulated maintenance state during torpor progression and then undergoes rapid molecular reactivation upon arousal.

The pathway analyses further support this interpretation. Genes contributing most strongly to PC1 were enriched not only for metabolic and redox-related functions, but also for RNA-processing pathways, particularly RNA splicing and ribonucleoprotein-related processes. This suggests that torpor progression is associated with coordinated regulation of RNA handling, proteostatic control, and metabolic adaptation in the cortex. This is consistent with the idea that cortical neuronal tissue must maintain essential protective functions while minimizing energy expenditure under conditions of strongly reduced metabolism. By contrast, genes contributing to PC2 were associated more strongly with intracellular signaling, protein trafficking, and regulatory processes, indicating that separation along this axis reflects the shift between torpor and arousal states. The enrichment of oppositely regulated genes between TE–TL and TL–AE, together with the associated KEGG pathways, further indicates that arousal is not simply a passive return to baseline but involves coordinated reversal of torpor-associated programs. In this context, disease-labeled KEGG categories do not reflect pathology but shared mitochondrial, metabolic, and stress-response modules that become engaged during the transition out of torpor.

These findings fit with the broader view that brain adaptation during hibernation is region-specific and highly dynamic. Previous transcriptomic studies have largely focused on the hypothalamus or whole brain, emphasizing pathways related to circadian control, metabolism, stress response, and seasonal regulation. Our results extend this literature by showing that the cerebral cortex itself displays a distinct and highly structured transcriptional response across torpor and arousal. This is important because the cortex must remain structurally viable through prolonged metabolic suppression and then rapidly recover function during arousal. The present data therefore suggest that cortical adaptation is not just a passive consequence of systemic hibernation, but reflects an actively regulated program of metabolic restraint, cellular maintenance, and reactivation. The strong reversal observed between late torpor and early arousal particularly supports the idea that the cortex is poised for rapid restoration of neuronal function when animals return to euthermia.

Several limitations should be considered. First, the sample sizes per physiological state were modest, which may reduce power for detecting small but biologically meaningful changes. Second, annotation remains incomplete in a non-model species, and some differentially expressed transcripts could therefore not be confidently assigned to known genes or pathways. Finally, the sampled states represent discrete snapshots of a continuous physiological cycle, meaning that rapid transitional events may not be fully captured. Future work using denser temporal sampling, cell-type-resolved approaches, and integration with proteomic or phosphoproteomic data will be important for defining how cortical regulation is coordinated across torpor and arousal at higher resolution.

Overall, our findings identify late torpor and especially early arousal as critical regulatory checkpoints in the garden dormouse cortex. The data support a model in which cortical adaptation to hibernation relies on coordinated metabolic suppression during torpor progression followed by active transcriptional reversal during arousal, thereby helping preserve neuronal integrity across repeated torpor–arousal cycles.

## Conclusion

This study shows that the cerebral cortex of the garden dormouse undergoes highly stage-specific transcriptional regulation across the hibernation cycle. The strongest transcriptomic remodeling occurs during progression into late torpor and, most prominently, during the transition from late torpor to early arousal, whereas entry into torpor and later arousal involve comparatively limited change. Functional analyses indicate that these transitions are associated with metabolic, stress-related, and regulatory programs consistent with reversible cellular maintenance and rapid functional recovery. Together, these findings identify the cortex as an actively regulated tissue during hibernation and highlight late torpor and early arousal as key phases for understanding how neuronal integrity is preserved under extreme metabolic suppression.

## Scripts availability

https://github.com/Deidarsik/Garden-dormouse-cerebral-cortex-git

